# Attentional Modulation of Macaque Visual Processing Areas

**DOI:** 10.1101/345439

**Authors:** Gaurav H. Patel, Lawrence H. Snyder, Maurizio Corbetta

## Introduction

The visual expectation that an object will appear at a certain location or time may increase the accuracy and speed of its detection (Eriksen and Hoffman, 1974; Posner, 1980; Carrasco et al., 2000; Dosher and Z-L., 2000). The ability to use prior information to enhance the processing of visual stimuli at a specific location is critical for quickly sorting through the myriad of incoming stimuli and making decisions regarding future actions. This selection process, known as visual attention, appears to be fundamental to our ability to navigate and interact with the environment, and accordingly has been the focus of innumerable psychological and neurobiological studies (Pashler, 1998).

Knowledge about the neurobiology of attention has traditionally come from neuropsychological and functional neuroimaging studies in human subjects and neurophysiological studies in monkeys (Kastner and Ungerleider, 2000; Corbetta and Shulman, 2002; Reynolds and Chelazzi, 2004). Studies in of macaque monkeys have been especially helpful in relating the neuronal mechanisms of attention to the underlying anatomy and physiology, which are much better understood for the monkey than the human brain. While the overall framework and principles underlying the neurobiology of attention are fairly consistent for humans and monkeys, the lack of a direct means of comparing the human fMRI and macaque electrophysiology data has left many uncertainties about the processes underlying visual attention in the two species. These uncertainties may largely be due to the use of different techniques for measuring neural activity.

It is generally agreed that visual attention operations involve the recruitment of a distributed network of areas: occipital areas involved in the analysis and identification of visual stimuli, and higher-order areas in prefrontal and posterior parietal cortex involved in encoding expectations, maintaining task goals, and controlling the locus of attention (Colby and Goldberg, 1999; Kastner and Ungerleider, 2000; Miller and Cohen, 2001; Corbetta and Shulman, 2002; Moore and Armstrong, 2003; Reynolds and Chelazzi, 2004). However, while the distributed nature of visual processing activations in humans was immediately apparent early on in neuroimaging studies, this distributed view in monkeys has been only inferred from recordings carried out serially across different cortical areas in different studies and usually different animals. As a result, the full complement of areas recruited during a controlled visual attention task in monkeys is still unknown, and in some cases even the borders of these areas are unclear.

Another area of relative disagreement is the source of top-down expectation signals in monkeys and humans. Some single unit studies in monkeys and neural theories of attentional control have emphasized prefrontal cortex (ventral and dorsal) as the source of top-down control (Desimone and Duncan, 1995; Miller and Cohen, 2001). Other studies have also implicated the frontal eye fields (FEF) and the lateral intraparietal area (LIP) in playing some role in either representing or controlling the locus of attention (Bisley and Goldberg, 2003; Moore et al., 2003; Moore and Fallah, 2004; Schall, 2004; Wardak et al., 2004; Thompson and Bichot, 2005; Wardak et al., 2006). Because of the problems in comparing studies outlined above, it is still unclear how these areas interact in the control of attention in the macaque. In contrast, neuroimaging studies in humans identified early on a dorsal fronto-parietal network including the putative human FEF and posterior parietal cortex as the most likely source of spatial and non-spatial biases to visual cortex (Corbetta and Shulman, 2002). Human prefrontal cortex, on the other hand, has mainly been implicated in working memory (McCarthy et al., 1994). No comparison of these two views of the neural systems of top-down control has been performed to date.

A final area of uncertainty concerns the effects of attention on visual responses in striate and extrastriate visual cortex. Single unit studies have reported modulations of neural activity related to spatial and feature attention in intermediate and high-level visual areas (e.g. V4, MT, IT, PPC) but much less robustly in low-level visual areas (e.g. V1, V2) (Motter, 1993, 1994; Luck et al., 1997; Chelazzi et al., 1998; Gottlieb et al., 1998; McAdams and Maunsell, 1999; Seidemann and Newsome, 1999; McAdams and Maunsell, 2000; McAdams and Reid, 2005). In fact, there have been only a handful of studies that have measured in the same animals attentional modulations of visual activity in different areas (Motter, 1994; Luck et al., 1997). In particular weak or no attentional modulations have been reported in area V1, primary visual cortex (but see McAdams)(Luck et al., 1997; McAdams and Reid, 2005). Conversely, human neuroimaging studies of visual attention modulations have provided a somewhat different picture. Stronger attentional modulations have been measured in intermediate and high level visual areas (V4, MT, IT) than in low-level visual ones (V1,V2) (Pinsk et al., 2004; Serences and Boynton, 2007). On the other hand, numerous experiments have reported robust and significant modulations related to spatial attention in human primary visual cortex (Gandhi et al., 1999; Pinsk et al., 2004; Serences and Boynton, 2007; Sylvester et al., 2007).

Electrophysiological analysis of these modulations have also suggested that V1 attentional modulations in humans reflect relative late feedback signals from higher-order regions rather than early bottom-up recruitment from the primary sensory volley of inputs (Martinez et al., 1999). Whether the difference in the degree of early visual cortex modulation, especially V1, reflects a methodology or species diffference remains to be seen (Posner and Gilbert, 1999). Current commentaries on this point tend to highlight the fact that neuroimaging measures of neuronal activity are sensitive to both spikes and total changes in local field potential, a measure that captures inputs and local processing within an area, whereas single unit recordings of spike rates are more biased toward the output of large pyramidal cells (Logothetis and Wandell, 2004). Therefore, if the V1 modulation does indeed reflect top-down feedback it may be in fact more evident in fMRI than neural measures.

To address these issues we measured neural activity with functional magnetic resonance imaging (fMRI) blood oxygen level dependent (BOLD) signals from two awake-behaving macaques performing a difficult visual attention task. The animals covertly monitored one of two peripheral RSVP streams while looking for a specific target object in a stream of difficult-to-discriminate distracter objects. To measure attentional modulations of stimulus-evoked activity in the whole brain we compared BOLD signals to one stream of stimuli when it was attended as compared to when it was unattended. Given only the side of attention was varied while the visual stimulation was kept constant, any change in activity can be interpreted as reflecting the interaction of an attentional set for the location and object of interest with the incoming stream of stimuli.

## Materials and Methods

### Animal Subjects, Surgery, and Experimental Setup

Two male monkeys (Y and Z, Macaca mulatta, 5-7 kg) were used in accordance with Washington University and NIH guidelines. Prior to training, surgery was performed in aseptic conditions under isofluorane anesthesia to implant a head restraint device. The head restraint was constructed from polyetheretherketone (PEEK) and was anchored to the skull with dental acrylic and 12-16 ceramic screws (4 mm diameter, Thomas Recording GmbH, Germany).

The monkeys were trained to perform the task in a setup that simulated the fMRI scanner environment. In both the training setup and scanner, the animal sat horizontally in a “sphinx” position inside of a cylindrical monkey chair with its head rigidly fixed by the restraint device to a head holder on the chair (Primatrix Inc., Melrose, MA). An LCD projector (BENQ, Irvine, CA) was used to present visual stimuli on a screen that was positioned at the end of the bore, 75 cm from the monkeys’ eyes. A flexible plastic waterspout was positioned near the animal’s mouth for delivery of liquid rewards.

Eye movements were monitored by an infrared tracking system (ISCAN Inc., Melrose, MA). A camera positioned outside of the bore monitored the left eye through a hole in the bottom of the screen; a plastic tube running from the camera to the hole prevented extraneous light from entering the bore via the hole. The eye was illuminated with an IR light source positioned 5-6 cm below the monkey’s left eye. The eye position coordinates were relayed to the behavioral control system, which consisted of two linked computers running custom software. Blinks were detected and the resulting artifacts in the eye-position signal were compensated for on-line.

The monkey was trained to place its paw inside a box that was affixed to the front of the chair under the monkey’s chin, and to rapidly withdraw and replace its paw when a target appeared. These movements were detected by a photoelectric sensor (Banner, Minneapolis, MN). The behavioral control system recorded eye position, hand status and scanner synchronization signals, presented visual stimuli, and delivered rewards. For further details about the experimental setup, see Baker *et al.* (Baker et al., 2006).

### Display and Task

At the beginning of a BOLD run, a white square (.3°x.3°) appeared at the center of the screen (see **Figure 1b**). To begin the fixation period, the monkey was given 2 seconds to position his eyes within 1.1° horizontally and 5° vertically of the fixation point, and then another 4 seconds to insert his hand into the hand-response device. The eye-window was longer in the vertical axis to accommodate residual artifacts in the eye-tracking signal stemming from both blinks and pupil constriction. A 15 second fixation period would then begin coincident with the start of the next MR frames (0-3s); at the end of 15 seconds the task block would begin. Two RSVP streams were started simultaneously, one at 5.3° eccentricity horizontally to the left of the fixation point, and the other at 5.3° to the right. The size of each object was scaled to 4.6° square, so that the images spanned 3°-7.6° visual eccentricity. The streams were on the screen for 60 seconds, followed by 30 seconds of fixation, and then the cycle repeated. The run continued until either the speed of the stream required adjustment or the monkey quit working.

**Figure 1.**
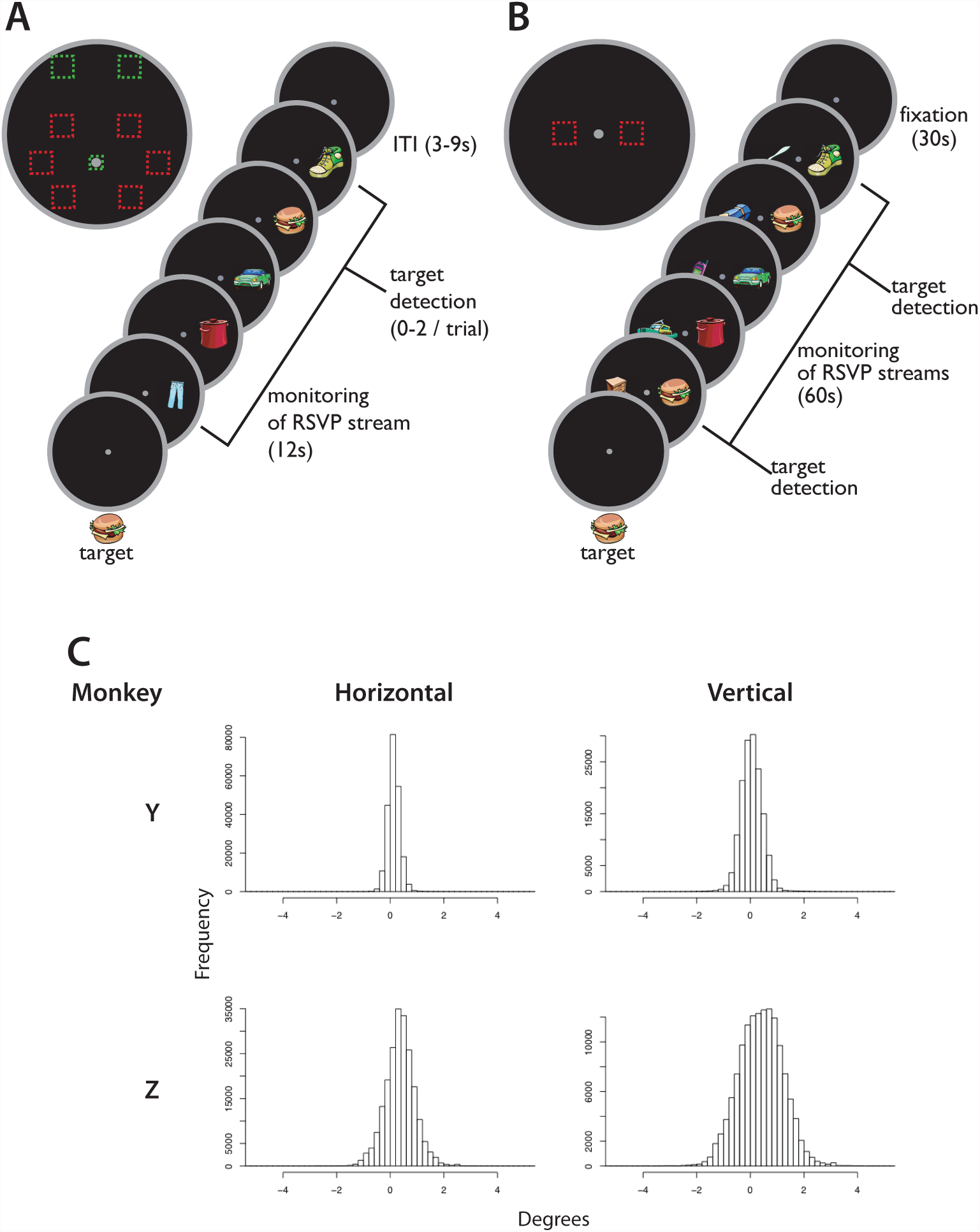
**a)** The single stream paradigm. **b)** The two-stream paradigm. **c)** The distribution of horizontal and vertical eye-positions throughout the single-stream polar angle task, sampled every 40ms. Positive values are in the direction of the presented stream.

The RSVP streams consisted of bitmap images drawn from a pool of 42 color illustrations of everyday objects (Barry’s Clipart Server, http://www.barrysclipart.com/). For each session, one of the 42 objects was selected as the target, and the 41 others left as distracters. The distracters were shown in random order with the constraint that the same image could not be shown twice in a row. For each BOLD run, we set the time that a single object was on the screen (stimulus duration) as short as possible to keep the monkey’s detection rate between 40% and 80%; for monkey Y the range of times was 112-250ms, and in monkey Z the range was 45-300ms. Each session, the target was chosen randomly from the 42 pictures, and one of the streams was randomly designated as the “target” stream. For that session, all of the targets appeared in the designated stream.

The monkey’s task was to detect the memorized target object while maintaining fixation on the central point. Targets appeared in the expected stream every 6 +/- 3 seconds. After the appearance of a target, the monkey had 1 second in which to indicate detection by moving his hand out of and back into the response box. Upon successful detection, the monkey was given a single reward. In addition, a reward was given after the presentation of 2-3 targets for the maintenance of task readiness (fixation and hand in the response device). During the fixation period, rewards were given once every 6 +/- 3 seconds for the maintenance of fixation. In monkey Z, an additional fixation reward was given every 6 seconds. In general, the number and relative sizes of rewards were adjusted to encourage fixation, maximize true hits and minimize false positives, and the size of the rewards was increased with the duration of the session.

If at any time during the run the monkey broke fixation, or if the monkey removed his hand from the device outside of the 1-second reaction time window (such as when falsely detecting a target), the monkey was penalized by blanking the display and aborting the trial. After a short pause, during which no stimuli were presented and no rewards could be obtained, the fixation point reappeared, and a new trial was begun. The duration of the pause varied from 2-9s between sessions—as short as possible but long enough to provide enough negative reinforcement to keep the monkey on task. In addition, if the monkey made several errors in a row, the trials were paused for 15-60 seconds, the length again depending on the monkey’s behavior. Failures to detect presented targets (misses) were not punished.

At the beginning of the sessions, a sequence of instruction trials lasting ~20 minutes acquainted the monkey with both the session’s target and which of the two streams he needed to attend to detect the targets, and also gave him practice discriminating the target from the distracters. The trials were also used to set the speed of the trials such that the monkey was detecting 40%-80% of the presented targets. Occasionally the previous session’s target as a distracter was eliminated for the rest of the session if the monkey kept mistakenly signaling detection for this object during the instruction trials.

### Data Collection

Functional and anatomical data were collected in a Siemens 3T Allegra MRI scanner (Siemens Medical Solutions, Erlangen, Germany). High-resolution structural images were collected in separate sessions, during which the animal was chemically restrained (10 mg/kg ketamine, 0.6 mg/kg xylazine, .05 mg/kg atropine). T1-weighted images were acquired using a magnetization-prepared rapid acquisition gradient-echo pulse sequence [MP-RAGE; (0.5 mm)^3^ isotropic resolution, flip angle = 7°, six acquisitions] and a volumetric transmit and receive coil (16 cm i.d.; Primatrix).

Functional data were collected using a gradient-echo echo-planar pulse sequence sensitive to BOLD contrast (T2*) (T2* evolution time = 25 ms, flip angle = 90°) and a transmit-receive surface coil (13 cm inner diameter; Primatrix). The coil fit around each animal’s head post and was saddle-shaped to provide more extensive brain coverage as compared to a planar surface coil. Fifty-two coronal slices, each with a square field of view (96 × 96 mm, 64 × 64 base resolution, dorsal-to-ventral phase-encoding) and a thickness of 1.5 mm, were obtained using contiguous, interleaved acquisition, and a volume repetition time (TR) of 3000 ms. This scanning protocol was chosen to cover the whole brain at an isotropic spatial resolution of (1.5 mm)^3^. The first four volumes of each run were excluded from the analyses to allow for the equilibration of longitudinal magnetization. A T2-weighted image was acquired at the beginning of one session using a turbo spin-echo sequence [TSE; (1×1×1.5mm)^3^ resolution, flip-angle = 150°, single acquisition).

In a data collection session, up to 1409 volumes (7 minutes) were collected in a single run; the scanner was only stopped if the stimulus parameters needed adjustment or if the monkey quit working. In monkey Y, a total of 5800 frames were collected in 11 runs in 9 scanning sessions, and in monkey Z a total of 7100 frames were collected in 8 runs in 8 scanning sessions. No runs/sessions were excluded due to excessive movement or poor performance.

### Preprocessing

In the polar angle and eccentricity experiments, all runs in which the monkey successfully completed more than 50% of the presented trials were included in the analysis (37/39 runs in monkey Y, 35/39 runs in monkey Z). The exception was monkey Y’s eccentricity data; this criterion was removed due to his tendency to increased errors resulting from quick out-and-back saccades to the peripheral streams. Because the error trials were modeled separately, the removal of this criterion did not appear to affect the data quality, as the statistical, focality of activations, and overall distribution of activity appeared to be similar in the two monkeys in this experiment. No runs were excluded for excessive movement. In the two-stream experiment no runs/sessions were excluded due to excessive movement or poor performance.

Each reconstructed fMRI run produced a 4-dimensional (x, y, z, time) data set that was passed through a sequence of unsupervised processing steps using in-house software. The data were first corrected for asynchronous slice acquisition using cubic spline interpolation, and also for odd-even slice intensity differences resulting from the interleaved acquisition of slices. Correction factors were then calculated for 1) normalization across runs, in which each four-dimensional data set was uniformly scaled to a whole brain mode value of 1000; and 2) a 6-parameter rigid body realignment to correct for within- and across-run movement. Finally, correction factors were calculated for aligning the data from each session first to each other and second to the monkey’s own T2-weighted image, aligned to the macaque F6 atlas (http://sumsdb.wustl.edu/sums/archivelist.do?archive_id=6636170) using a 12-parameter affine registration algorithm (Snyder, 1995). All of these correction factors were then applied to the data in a single resampling step. To further refine the cross-session alignment within each monkey, an additional recursive alignment algorithm was implemented. Atlas-aligned representative images from each session were averaged together, and then each session’s representative image was realigned to this average using the 12-parameter affine registration. These newly aligned images were then averaged together and realigned to the T2-weighted image, which in turn was used as the target for the next realignment iteration. This was repeated until the change in variance averaged across all the voxels across all of the sessions asymptotically reached a minumum. Finally, the final atlas-registration matrix and the above-calculated normalization and movement-corrections factors were re-applied to the data in a single resampling step.

### Projection of the statistical data to the cortical surface

For each monkey, a cortical surface model was created by segmenting the gray and white matter of the monkey’s own MPRAGE (http://brainvis.wustl.edu, (Van Essen, 2002)). The resulting segmentation volume was then edited to fit the average atlas-aligned EPI image that resulted from the above-described atlas alignment scheme. The cortical surface was then flattened and registered to the macaqueF6 surface atlas (http://sumsdb.wustl.edu/sums/archivelist.do?archive_id=6636170) using sulcal-folding markers. To project statistical maps (see below) to the flattened, registered cortical surface, the volume maps were resampled to (.5mm)^3^, and then each point on the surface was painted with the highest absolute z-score within 1.5 mm. For comparisons between the four hemispheres, these maps were then warped to the macaqueF6 atlas right hemisphere surface by way of the sulcal-folding surface registration procedure described above.

### Analysis of Attentional Modulation

Statistical maps of activity were created using a general linear model (GLM) implemented with in-house software. To create z-statistic maps, each event-type was modeled as an independent regressor formed by convolving a gamma-function (Boynton et al., 1996) with a boxcar of a specified duration. The included regressors were the 60 second streams (separated by whether the session was “attend-left” or “attend right”), streams that were cut short by extended punishes, the target detections (.5 seconds), bonus rewards (.5 seconds), and missed targets (.5 seconds). Additional regressors were included for the fixation-break and false-detection error pauses, and also linear trend and baseline. If the punish period lasted longer than 21 seconds, those frames were excluded from the analysis. In the model, only the amplitude of the regressors was allowed to vary freely for each voxel, and the standard error was estimated from the remaining variance. From these amplitudes and standard errors, voxel-wise z-statistic map contrasting each condition versus baseline was created.

The “attend-left” and “attend-right” contrast maps were summed together to give a map of voxels stimulated by visual stimuli, which was then used to create regions-of-interest (ROIs) from significantly activated clusters of voxels. The ROIs were created in volume space in each monkey using an automated peak-search algorithm that grouped together voxels within 6mm of a local maxima. This procedure, however, sometimes arbitrarily divided clearly contiguous foci of activity. To remedy this, the various atlases available in Caret and the Saleem and Logothetis atlas were used to combine and label these automated ROIs. In general, unless clear anatomical criteria could be used to divide neighboring ROIs into separate clusters, adjacent clusters were combined to form a single ROI. This procedure gave us four occipital (V1, V2/V3d, V2/V3v, V4) and seven extra-occipital (LIP, MT, PITd, CITd, AITd, FEF, and area 46p) ROIs for each hemisphere. The lone exception was the absence of a CITd in monkey Z’s right hemisphere. These ROIs were then used to extract the magnitude of activation evoked by the streams in the “attend-left” and the “attend-right” conditions.

For the ROIs in each hemisphere, these magnitudes were reorganized into “attend-contralateral” and “attend-ipsilateral” categories, and the difference in magnitude (contra-ipsi) was calculated. Then for each ROI, a paired t-test was used to compare the magnitudes for the two conditions across the four hemispheres. Also the difference between the two conditions was averaged for each ROI across hemispheres and used to calculate a percent-modulation ((contra-ipsi)/ipsi). To determine the significance of the difference between the two attention conditions for each ROI in each hemisphere, we calculated the z-score of the difference between the amplitudes of the regressor fit to the response for each condition averaged across the entire ROI. To verify the veracity of the magnitude estimates, the ROIs were used to extract time-sources of the BOLD signal evoked by the attend-left and attend-right conditions for each ROI from a GLM in which the shape of the hemodynamic response function was not assumed.

### Eye position analysis

Although large eye position windows were enforced during data collection, actual eye position was generally very close to the fixation point. In order to quantify this, we plotted histograms of the horizontal component of the monkey’s eye-position calculated in 40ms bins. The components were plotted as the horizontal displacement towards the attended stream. The resulting eye-position distributions were used to calculate the mean and standard deviation of the mean horizontal eye-position for each monkey in the two-stream experiment.

## Results

We measured BOLD signals in two monkeys while they searched through one of two RSVP streams for a previously memorized target (see **Figure 1b**). The streams appeared simultaneously along the horizontal meridian, 5.3° to the right or left of a central fixation point for 60 seconds, followed by 30 seconds of fixation. To encourage the animal to covertly monitor just one stream for the duration of a session, all the targets appeared in that one stream. The stream containing all the targets was assigned randomly each session.

### Behavior

The presentation time of each stimulus within the stream was adjusted in each session to maintain the difficulty of the task. In monkey Y, the streams were varied in speed between 110 and 250 ms to maintain an average session detection rate of 53% (sd=18%); in monkey Z, the speeds were varied between 45 and 300ms, with a detection rate of 72% (23%). Mean reaction times for monkeys Y and Z were 450ms (46ms) and 428ms (28ms), respectively.

The two monkeys broke fixation less than once per block (monkey Y: 27 fixation breaks/182 blocks; monkey Z: 174/234); these fixation breaks were followed by blanking the display for 2-9 seconds and were modeled as regressors of no interest. The eye-position data during and immediately preceding these fixation breaks were excluded from further analysis. The average horizontal fixation error was .16° (sd=.26°) in the direction of the attended stream for monkey Y and .09° (.36°) in the direction of the attended stream for monkey Z (see **Figure 2a**). These values indicate a highly significant but very small bias in the direction of the attended stream (p<.0001). In fewer than .006% (Monkey Y) and fewer than .03% (Monkey Z) of samples were the eyes more than 2.5 deg from the fixation point. While the horizontal fixation error for both monkeys was significant, it was on average less than a 1/10^th^ of the distance between the fixation point and the inner edge of the RSVP stream and should not be able to explain the following results.

**Figure 2.**
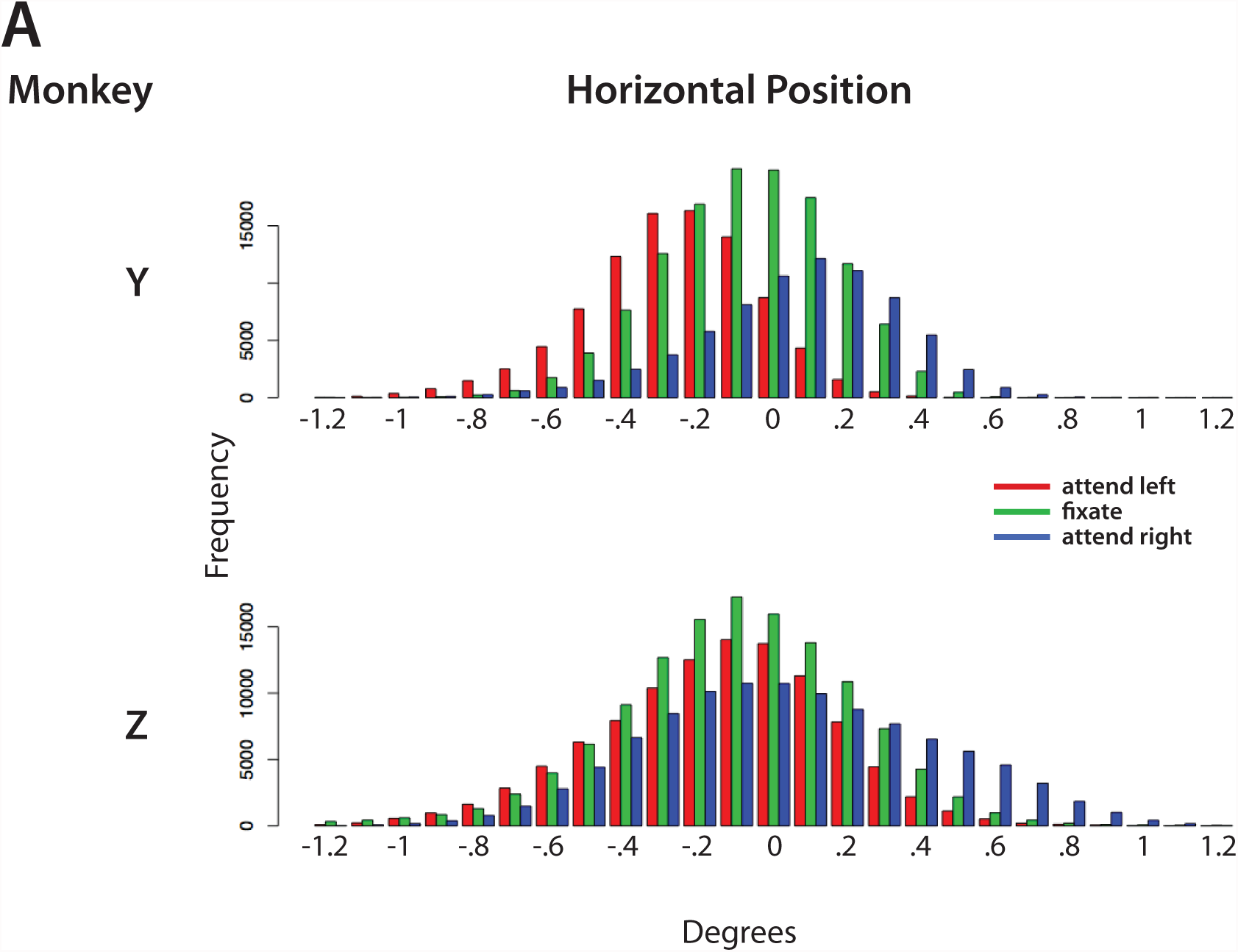
Horizontal fixation control during the two-stream experiments, sampled every 40ms. Negative values represent deviations to the left of the fixation point, and positive to the right.

### Areas involved in visual processing

The presence of the two RSVP streams on the screen evoked BOLD signals at locations throughout the cortex (see **Figure 3**). These included foci in visual cortex, inferotemporal cortex, posterior parietal cortex, and prefrontal cortex. In general, more voxels in the statistical maps in monkey Y passed the significance criterion than in monkey Z; this increased power in monkey Y resulted in a greater number of activity foci. However, the strongest foci generally matched in the two monkeys, and so only the foci consistent in both animals were considered further.

**Figure 3.**
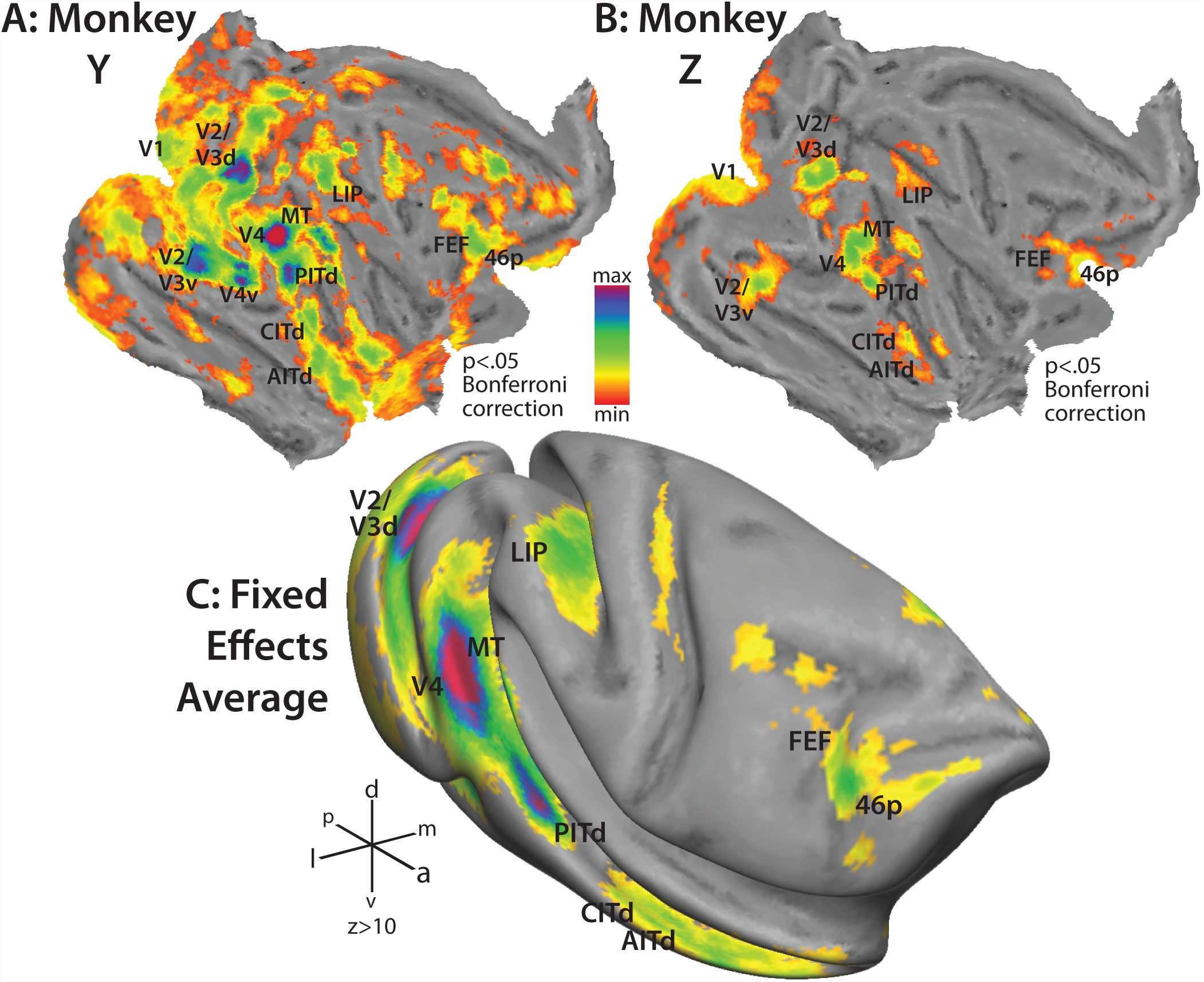
The cortical distribution of activity evoked by the two-stream paradigm. **a)** & **b)** Activity averaged across the two hemispheres in each monkey and projected to the flattened macaqueF6 surface. c) The fixed-effect average of the two monkeys projected to the macaqueF6 inflated surface.

In visual cortex, foci were seen in the caudal portion of the calcarine sulcus, the posterior bank of the lunate sulcus, in the fundus of the inferior occipital sulcus, and on the gyral surface between the lunate and superior temporal sulci. These corresponded in location to the horizontal meridian representations in V1, between V2d and V3d, between V2v and V3v/VP, and V4, respectively. In monkey Y, an additional strong focus was seen at the ventral anterior border of V4 (labeled V4v in the monkey Y flat-map in **Figure 3a**) but did not appear in monkey Z; this discrepancy was most likely due to differences in the susceptibility artifact stemming from the nearby ear canals.

In the IPS, the focus of activity spanned much of the lateral bank of the IPS, from the fundus to the gyrus; and starting at the bend of the IPS stretched ~1 cm in the axis parallel to the fundus. This focus covered much of areas LIPd and LIPv in the Lewis and Van Essen atlas. In monkey Y, another focus of activity was observed on the lateral bank at the junction of the parietal-occipital and intraparietal sulci, but was absent in monkey Z not considered further. The flat projection shows activity on the medial bank as well as on the lateral bank (see monkey Z in **Figure 3b**), but inspection of the volume data shows that the activity is in fact confined to the lateral bank, so that the medial bank activation in the flat map is a misprojection of the volume-space statistical data.

In the superior temporal sulcus, the most caudal focus of activity was immediately adjacent to V4. The peak voxels fell near the border of V4tp/a and MT in the Lewis and Van Essen atlas, so the ROI was labeled MT. The MT focus stretched about 7mm along the lateral-inferior bank of the caudal portion of the STS. 7mm rostral to MT was another large focus of activity. The boundaries of this focus fit those of PITd in the Felleman and Van Essen atlas, and also within the larger borders of TEO as delineated by Ungerleider and Desimone (Desimone and Ungerleider, 1986a; Ungerleider and Desimone, 1986; Felleman and Van Essen, 1991; Van Essen, 2002; Van Essen, 2003). PITd was about 5mm long in the rostral-caudal axis, and on the lateral portion of the inferior bank of the STS, near the lip of the gyrus. 5mm rostral to PITd in 3 of the 4 hemispheres was a pair of foci of activity (in the 4^th^ hemisphere there was only one focus 10mm roistral to PITd). These two regions constitute CITd and AITd in the Felleman and Van Essen atlas, and still fall within area TEc in the Ungerleider and Desimone partitioning scheme. Both of these foci also fell along the lateral portion of the inferior bank of the STS, and the activity in monkey Y extended rostrally nearly to the temporal pole. The activations on the superior bank of the STS in **Figure 3** are misprojections from the inferior bank.

In prefrontal cortex, two main foci of activity were observed. The first was along the anterior bank of the arcuate sulcus, stretching from the midpoint 4mm laterally into the inferior ramus. This focus was split between 8Ac and 6Vam in the Lewis and Van Essen atlas and corresponded to FEF in the Saleem and Logothetis atlas (Lewis and Van Essen, 2000b; Saleem and Logothetis, 2006). Anterior to FEF was the other focus, which covered much of the cortex on the gyrus anterior to the arcuate sulcus and continued into the posterior portion of the principal sulcus. This focus corresponded to area 46p in both the Lewis and Van Essen and Saleem and Logothetis atlases. Additional foci were observed in both monkeys near the inferior termination of the arcuate sulcus; these foci were strong in monkey Y but weak in monkey Z and not considered further.

### Modulation of Visual Stimulus Processing

To see the effects of attention on visual stimulus processing, we calculated an average magnitude of the evoked BOLD signal for distinct regions in each hemisphere for both the attend-contralateral and attend-ipsilateral conditions (see Methods for details of ROI determination). For each ROI the magnitudes were averaged across the four hemispheres for each condition. Then we calculated both the mean difference between attend-contralateral and attend-ipsilateral conditions (see **Figure 4** and **Table 1**) and a percent modulation (contra-ipsi/ipsi).

**Figure 4.**
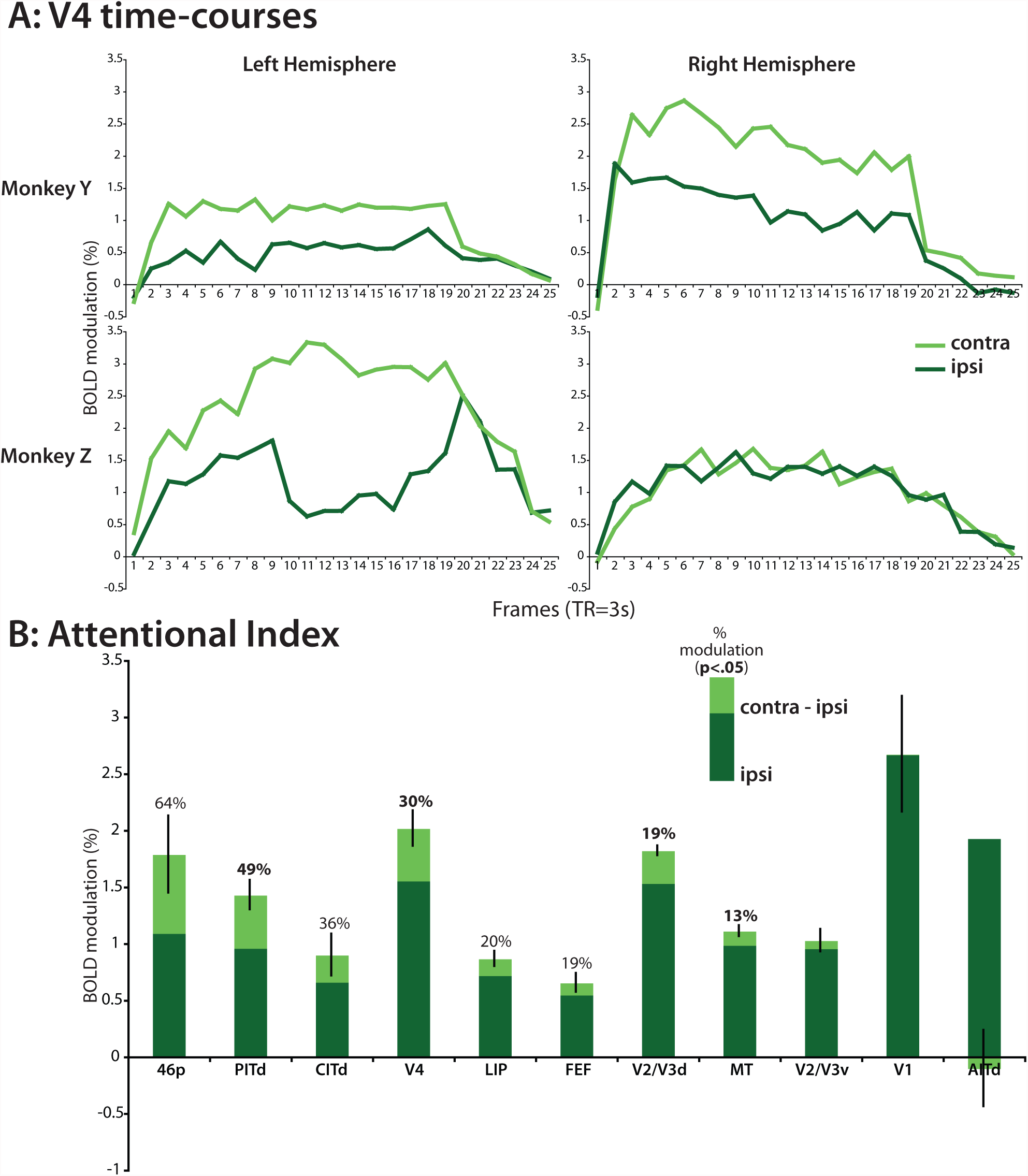
Attentional modulation in visual processing areas. **a)** Example time-courses from V4. **b)** Magnitude of modulation in all of the visual processing areas. Percentage reflects the amount of modulation over the ipsilateral magnitude (contra-ipsi/ipsi). Bolded percentages indicate sginificance of p<.05; non-bolded percentages are p<.2.

**Table 1.**
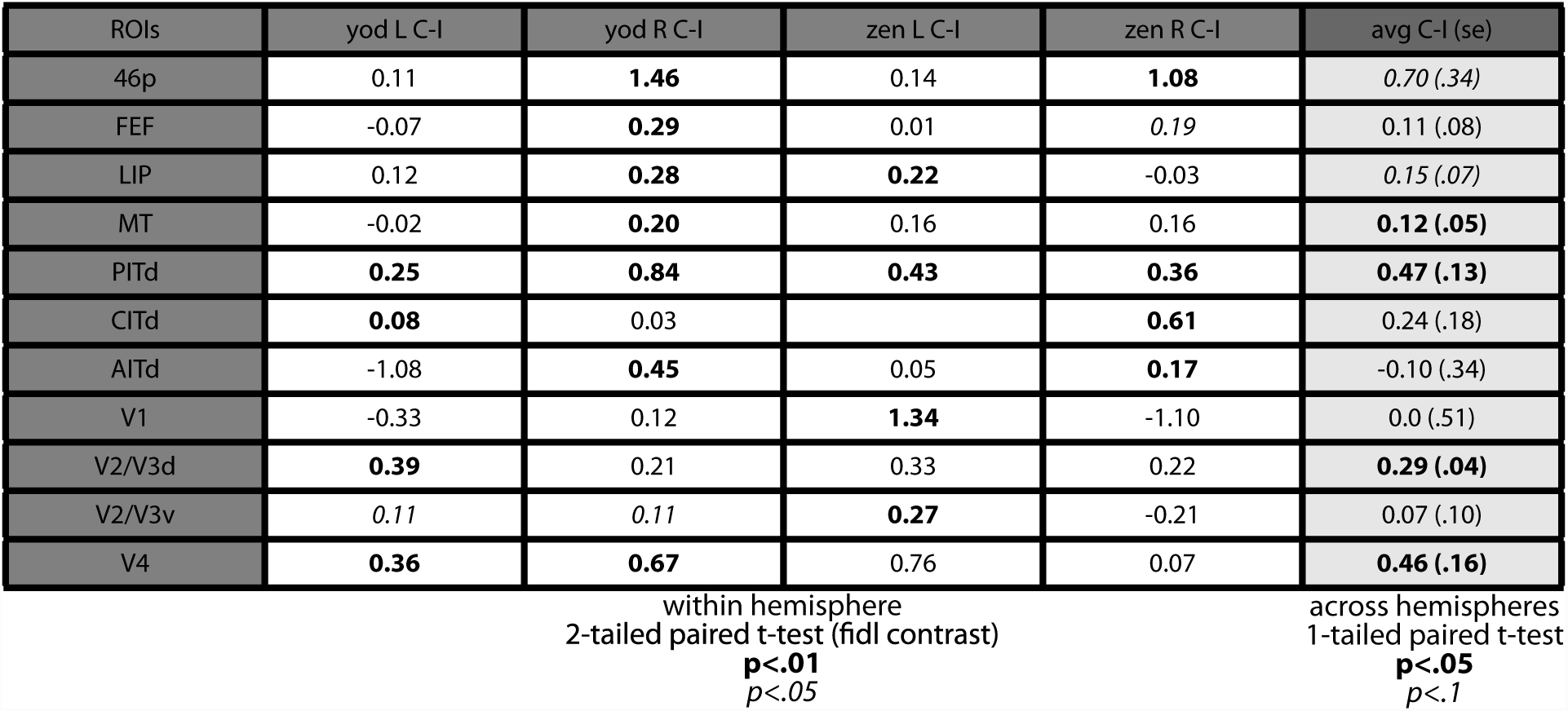
The magnitude of modulation in each ROI in each hemisphere. The given magnitudes are the difference in BOLD modulation between the attend-contralateral and attend-ipsilateral conditions. For the individual hemispheres, bolded values indicate significance at p<.01, and italics indicate p<.05. For the averages, bold indicates p<.05 and italics p<.1.

**Figure 4b** displays the modulations in order from the strongest to the weakest. The strongest modulations were observed in area 46p (64%, p=.067, one-tailed paired t-test), but the effect of attention appeared to be lateralized to the right hemisphere in both monkeys (monkey Y/Z: left hemisphere—18/10%, right hemisphere—118/90%). The next largest modulations were observed in the inferotemporal cortex area PITd, which demonstrated large (49%) and consistent effects of attention, both at the population level (n=4 hemispheres, p<.05, one-tailed paired t-test) as well as in each individual hemisphere (p<.05, two-tailed t-test). PITd was followed by the more rostral inferotemporal area CITd (36%); however, CITd could be identified in only three of the hemispheres, and at the population level only presented a weak trend (p=.16). V4 was next, with an average modulation of 30% across the four hemispheres (p<.05). In LIP and FEF, the magnitude of the effect of attention was weaker (20% in LIP, 19% in FEF), but only showed a strong trend in LIP (p=.059, one-tailed paired t-test) and a weaker trend in FEF (p=.14). The smallest effects were observed mostly in visual cortex, with small but consistent modulations in V2/3d (19%, p<.05) and MT (13%, p<.05), and then insignificant and near-zero modulations in V2/3v and V1. The inferotemporal cortex area AITd also demonstrated insignificant effects of attention. The relative sizes of the modulations demonstrate that in general higher-level visual areas exhibited greater effects of attention than did most of the occipital areas.

## Discussion

By using BOLD-fMRI to assess the level of neural activity across the entire cortex in two macaques monitoring peripheral RSVP streams for the appearance of a target, the results of this study define the distributed network of visual processing areas involved in visuospatial attention. We have also found that the modulation of the BOLD signals by attention matched those observed in previous human and macaque studies; furthermore, the differences in these magnitudes across the network of areas may indicate differences in the roles each of these areas play in visual attention. Lastly, we found support for potential differences between humans and macaques in the effect of attention on early visual areas.

The areas involved in visual processing in our study include those in frontoparietal and inferotemporal cortex, and early visual areas up to but not including V1. This distribution of areas matches the network of visual processing areas as delineated by connectional anatomy studies of V4, PITd/TEO, FEF, and LIP (Andersen et al., 1990; Felleman and Van Essen, 1991; Distler et al., 1993; Schall et al., 1995; Lewis and Van Essen, 2000a; Ungerleider et al., 2007). Chemical deactivations and lesions of these visual processing areas lead to impairments in the discrimination of visual stimuli or in the allocation of visual attention (De Weerd et al., 1999; De Weerd et al., 2003; Wardak et al., 2004; Buffalo et al., 2005; Wardak et al., 2006; Rossi et al., 2007).

While inferotemporal cortex is known to be involved in the discrimination and identification of visual objects, there is little agreement about the partitioning of this large region (Felleman and Van Essen, 1991; Distler et al., 1993; Lewis and Van Essen, 2000a; Van Essen, 2003). The results of the present study support a hybrid of the available layouts. At roughly the midpoint of the rostral-caudal axis of the STS, we found an area strongly activated by the visual processing task and modulated by the direction of attention. This focus of activity corresponded to the area PITd as defined by both the Felleman and Van Essen and Distler *et al.* partitioning schemes (Felleman and Van Essen, 1991; Distler et al., 1993). These studies separated PITd inside the STS from the more lateral/ventral TEO, which contained a topographic map of the periphery on the ventral surface of the temporal lobe. Rostral to PITd we found two foci that were not strongly modulated by the direction of attention. While these two foci were encompassed by the larger area labeled TEa in the Lewis and Van Essen and Distler partitioning schemes, they appeared to be distinct foci—as evidenced by the lack of only one of these areas in monkey Z—which fit the Felleman and Van Essen definitions of AITd and CITd (Felleman and Van Essen, 1991). In monkey Y, there were additional foci of activity within the bounds of TEa, suggesting that further divisions of this area may exist.

Prefrontal cortex is another region in which the partitioning scheme has not been clear, and is often treated as a mixed bag of functions. We observed two distinct foci, one in FEF and the other in the ventral portion of area 46p as defined by the Lewis and Van Essen atlas; there was no other consistent foci of activity outside of the ventral portion of area 46p (Lewis and Van Essen, 2000b). The ventral portion of the focus matched with the recording sites of studies of working memory in object discrimination tasks (Wilson et al., 1993; Miller et al., 1996) and the dorsal portion matched the posterior parts of areas implicated in spatial processing (Wilson et al., 1993). There was some evidence that the area 46p activation consisted of two foci split dorsal-ventral as outlined by Wilson *et al.*, but the separation was not consistent across hemispheres. In general, these results provide additional constraints for previously detailed partitioning schemes by demonstrating that clear functional boundaries exist in prefrontal cortex.

By using BOLD-fMRI to assess activity across the entire brain at once, we are also able to compare activity levels across all of the visual processing areas. In general, the amount of modulation (10-50%) appeared to be roughly agree with results from macaque electrophysiology results (Miller et al., 1996; Luck et al., 1997; McAdams and Maunsell, 1999; Seidemann and Newsome, 1999; DeSouza and Everling, 2004; Sereno and Amador, 2006). Note that while we have treated the modulations as increases in activity related to the processing of attended stimuli, the effect is likely to be a combination of this increase and the suppression of the processing of the ipsilateral stream (Reynolds and Chelazzi, 2004). However, because of the whole-brain coverage of the BOLD-fMRI technique, we were also able to see systematic differences in the amount of modulation in the different areas: inferotemporal and prefrontal areas tended to exhibit large effects of attention, frontoparietal areas demonstrated intermediate effects, and occipital visual areas demonstrated intermediate or no effects. We observed two trends in this data.

The first trend was that in the areas falling within the inferotemporal areas (PITd, CITd, and AITd), the further rostral an area was, the less the effect of attention. This matches the increase in receptive field size seen in traveling caudal to rostral through the inferotemporal cortex (Desimone et al., 1984; Desimone and Ungerleider, 1986b; Boussaoud et al., 1991). In the most rostral temporal areas, the receptive fields can extend into the ipsilateral field. This trend in receptive field sized may mean that attention to either stream enhanced the activity in these areas, which would lead to decreasing differences in attending contralaterally versus ipsilaterally with increasing receptive field size.

Second, areas previously implicated in the discrimination of the features of stimuli appeared to demonstrate the largest effects, namely V4, PITd, CITd, and 46p (Ungerleider et al., 1993; Chelazzi and Desimone, 1994; Miller et al., 1996; De Weerd et al., 1999; Chelazzi et al., 2001; De Weerd et al., 2003; Buffalo et al., 2005). LIP and FEF exhibited smaller effects, possibly suggesting that, in line with traditional views and contrary to recent reports, much of feature processing does take place in inferotemporal and not parietal cortex (Sereno et al., 2002; Denys et al., 2004a; Sereno and Amador, 2006). MT also demonstrated small effects of attention, in keeping with its apparent role in motion and not object discrimination (Maunsell and Newsome, 1987; Newsome and Pare, 1988; Seidemann and Newsome, 1999). Earlier visual areas also demonstrated small or no effects of attention, in agreement with previous electrophysiology studies (Motter, 1993; Luck et al., 1997).

In general, the results indicate that the effect of attention is strongest in the regions most critical in performing the necessary operations to discriminate the stimuli in this task. A series of lesion studies found that areas TEO (including PITd) and V4 were necessary for monkeys to filter stimuli and for the TE neurons in these monkeys (inclusive of CITd and AITd in our monkeys) to demonstrate effects of the distracters on the activity related to the processing of a target (Ungerleider et al., 1993; De Weerd et al., 1999; De Weerd et al., 2003; Buffalo et al., 2005). Whether the effects of attention on each of these visual processing areas would change with a difference task will require additional attention experiments with different types of stimuli.

Similar results have been observed in human fMRI studies. The regions involved in visual processing tend to be in the human homologues of the macaque regions described above, and magnitude of the modulations in these regions tends to be 10-50%, just as in the macaque (Gandhi et al., 1999; Pinsk et al., 2004; Sylvester et al., 2007). A study from Hemond *et al.* of object-discrimination also finds declining contralateral bias in higher-level ventral stream areas, and a recent study from Serences *et al.* finds that the IPS and human FEF regions demonstrate less contralateral bias than human V4 (Hemond et al., 2007; Serences and Boynton, 2007).

However, many studies of visual attention in the human also report robust attentional modulations in V1 (Gandhi et al., 1999; Kanwisher and Wojciulik, 2000; Pessoa et al., 2003; Bahrami et al., 2007; Sylvester et al., 2007; but see Yoshor et al., 2007). These modulations can be as much as 40% over the “attend-away” condition (Gandhi et al., 1999). Similar effects have been searched for in macaque electrophysiology studies, but the modulations are often absent or subtle as compared to the human fMRI results (Motter, 1993; Luck et al., 1997; Mehta et al., 2000).

Our results indicate the macaque V1 is poorly modulated by spatial attention even when these signals are measured with fMRI. Therefore the reported discrepancy between human fMRI and monkey single unit results does not reflect differences in the type of neural activity sampled (Logothetis and Wandell, 2004). Rather, it may represent a real interspecies difference in the effect of attention on V1 neural activity. Other fMRI studies of humans and macaques have detailed potential differences in higher-level visual processing areas (Denys et al., 2004a; Denys et al., 2004b; Koyama et al., 2004; Orban et al., 2004); these results indicate that species differences may extend all the way to primary sensory areas, where homologies were previously thought to be strong between the two species. While it is too early to tell whether these functional-anatomical differences translate to differences in the underlying attentional mechanisms, our results highlight the need to be cautious when applying macaque data to human models of cognition.

